# The regulation of methylation on the Z chromosome and the identification of multiple novel Male Hyper-Methylated regions in the chicken

**DOI:** 10.1101/2023.03.27.534313

**Authors:** A. Höglund, R. Henriksen, A.M. Churcher, C.G. Guerrero-Bosagna, A. Martinez-Barrio, M. Johnsson, P. Jensen, D. Wright

**Affiliations:** AVIAN Behavioural Genomics and Physiology Group, Linköping University, Linköping 58183, Sweden; NBIS, Umeå University, Department of Molecular Biology, 901 87, Umeå Sweden; NBIS, Uppsala University, SciLifeLab, Box 596, S-75124 Uppsala, Sweden; Department of Animal Breeding and Genetics, Swedish University of Agricultural Sciences, Box 7023, 750 07 Uppsala, Sweden

## Abstract

DNA methylation is a key regulator of eukaryote genomes, and is of particular relevance in the regulation of gene expression on the sex chromosomes, with a key role in dosage compensation in mammalian XY systems. In the case of birds, dosage compensation is largely absent, with it being restricted to two small Male Hyper-Methylated (MHM) regions on the Z chromosome. To investigate how variation in DNA methylation is regulated on the Z chromosome we utilised a wild x domestic advanced intercross in the chicken, with both hypothalamic methylomes and transcriptomes assayed in 124 individuals. The relatively large numbers of individuals allowed us to identify additional genomic MHM regions on the Z chromosome that were significantly differentially methylated between the sexes. These regions appear to down-regulate local gene expression in males, but not remove it entirely (unlike the lncRNAs identified in the initial MHM regions). In addition, trans effect hotspots were also identified that were based on the autosomes but affected the Z, and also that were based on the Z chromosome but that affected autosomal DNA methylation regulation. In addition, quantitative trait loci (QTL) that regulate variation in methylation on the Z chromosome, and those loci that regulate methylation on the autosomes that derive from the Z chromosome were mapped. Trans-effect hotspots were also identified that were based on the autosomes but affected the Z, and also one that was based on the Z chromosome but that affected both autosomal and sex chromosome DNA methylation regulation. Our results highlight how additional MHM regions are actually present on the Z chromosome, and they appear to have smaller-scale effects on gene expression in males. Quantitative variation in methylation is also regulated both from the autosomes to the Z chromosome, and from the Z chromosome to the autosomes.

## Introduction

DNA methylation is one of the key regulators of eukaryotic genomes, and can both inhibit (Gaston and Fried 1995, Mann, Chatterjee et al. 2013) and enhance gene expression (Yin, Morgunova et al. 2017, Höglund, Henriksen et al. 2020), depending on where the DNA methylation occurs. This DNA methylation can be environmentally driven (Nalvarte, Rüegg et al. 2018), but can also be modified and regulated via DNA variation (Kasowski, Kyriazopoulou-Panagiotopoulou et al. 2013, Kilpinen, Waszak et al. 2013, McVicker, van de Geijn et al. 2013, Bélteky, Agnvall et al. 2018, Guerrero-Bosagna 2019). We have previously addressed this using a wild x domestic chicken model to study the regulation of variation in autosomal DNA methylation, and how it can quantitatively regulate gene expression using a QTL mapping based approach. This enabled us to identify how domestication in the chicken led to a small number of large-effect trans hotspots, where these loci regulated variation in DNA methylation throughout the genome. Moreover, we found methylation to not only be the driver but also the response to gene expression variation (Höglund, Henriksen et al. 2020). However, the corresponding regulation of DNA methylation variation on the Z chromosome is still lacking. For example, the extent to which quantitative variation in DNA methylation is controlled between the autosomes and sex chromosomes is an open question, as is the extent to which DNA methylation is regulated on the Z chromosome in general. Given the role that DNA methylation plays in dosage compensation on the Z chromosome in the chicken, this is particularly relevant.

Dosage compensation prevents the expression imbalance originating from the number of sex chromosomes present in males or females when homo- and heterogametic sexes exist. Dosage compensation occurs when the dose effect due to one sex having only a single sex chromosome, and therefore half the number of gene copies, is compensated by either decreasing gene expression in the homogamete or increasing expression in the heterogamete. This can be over the whole sex chromosome or over specific regions (Mank 2013). Dosage compensation is less well-described in ZW systems, with the female typically being heterogametic, in contrast to the mammalian XY systems (Itoh, Melamed et al. 2007, Vicoso and Bachtrog 2011). In the mammalian XY system, dosage compensation is achieved by X inactivation, achieved via epigenetic mechanisms, notably DNA methylation and histone modification (Fang, Disteche et al. 2019). However, such chromosomal inactivation is largely absent from birds, with instead very incomplete and location-specific dosage compensation, if any (Ellegren, Hultin-Rosenberg et al. 2007, Mank and Ellegren 2009). Despite this, gene expression on the Z in males is not double that of females, but instead genes on the Z are on average around 30% upregulated in males (Ellegren, Hultin-Rosenberg et al. 2007, Melamed and Arnold 2007, Mank and Ellegren 2009).

DNA methylation still plays an important role for sex difference regulation on the avian Z chromosome. In particular, the Male Hyper-Methylated (MHM) region at 27.3Mb was first discovered by Teranishi and colleagues (Teranishi, Shimada et al. 2001), whilst more recently an additional region on 73.16-73.17Mb was also identified on the Z chromosome (Sun, Maney et al. 2019). With the initial region, it was found that males had greatly increased methylation in an approximately 500kb area, with nine genes that were present there not being expressed in males. In the case of the more recently discovered MHM region at 73.16Mb (designated MHM2), this was smaller and contained three long non coding RNAs (lncRNAs) that were female-biased in expression. In general, these studies are based on small numbers of samples, generally focussing on between species comparisons (for example, one great tit sample was used in Laine et al. 2016, two pooled samples from Whole Genome Bisulfite sequenced chicken were used in (Zhang, Yan et al. 2017), and one male and one female White Throated Swallow was used in (Sun, Maney et al. 2019)). This makes it harder to detect smaller regions, and in particular the scope of inter-individual variation in these MHM regions. This is concerning, particularly considering the degree of DNA methylation variation across individuals in populations and the role of methylation in phenotype formation (Heyn, Moran et al. 2013). Large-scale analysis of within species variation could give a better resolution of hypermethylated regions as well as detect differences between individuals in sex-specific methylation and gene regulation. Various questions still remain regarding the MHM regions, and the genes they contain. The sizes of the MHM regions and the effects of the decreased gene expression is particularly noteworthy – are these genes involved in fundamental sex differences? Similarly, are the genes within these MHM regions regulated in a region-by-region basis or on a gene-by-gene basis? Gene expression regulation via methylation is not restricted to solely promoter regions (Kasowski, Kyriazopoulou-Panagiotopoulou et al. 2013), but can affect gene expression (both positively and negatively) due to effects at enhancer sites, Transcriptional Elements (TEs) and the like. For example, our previous study based on autosomal methylation variation in the chicken found that there was a bias to being positively correlated, whilst correlations between methylation and gene expression could be found within a megabase upstream and downstream of the gene itself (Höglund, Henriksen et al. 2020). Given this, how far away from these MHM regions are genes being affected? Is this still affecting dosage compensation if it upregulates genes?

To investigate how DNA methylation variation is regulated on the Z chromosome, as well as the potential role of methylation in dosage compensation and sex differences, we conducted a DNA methylation quantitative trait locus (methylation QTL) analysis using an advanced intercross between domestic chickens and wild Red Junglefowl. We assayed the hypothalamic transcriptome and methylome on the Z chromosome for 124 individuals, having previously assayed the autosomes for these individuals. It was therefore possible to map both *cis* and *trans* related loci that modulate variation in DNA methylation on the Z chromosome, as well as to assess how methylation is used to regulate sex-differences in gene expression on the Z chromosome.

## Methods

The study population was composed of 124 chickens (55 females, 69 males) from which the hypothalamus tissue was dissected out at day 212. The individuals used were from an 8th generation advanced intercross, founded using a Red Junglefowl (wild) male and three White Leghorn (domestic) females. A detailed description of the intercross generation, housing conditions, etc can be found in (Johnsson, Gustafson et al. 2012).

### RNA and DNA methylation isolation

RNA was isolated from the hypothalamus tissue which was homogenised using Ambion TRI Reagent (Life Technologies) following the manufacturer’s protocol. cDNA synthesis and microarray-based gene expression were performed using a Nimblegen 135k array, as described previously (Johnsson, Williams et al. 2016). DNA was isolated from the remainder of the TRI reagent homogenate by mixing 125µl ice-cold 99% ethanol with 250µl TRI reagent homogenate. Samples were vortexed, incubated on ice for 5min and centrifuged at 12’000 RPM for 10 min in room temperature. The pellet was saved and isolation continued using the DNeasy Blood & Tissue Kit (Qiagen) following the manufacturer’s protocol. DNA methylation was assessed by Methylated DNA immunoprecipitation (MeDIP) protocol. Further details of the MeDIP protocol can be found in (Höglund, Henriksen et al. 2020).

### Phenotypes: methylation and gene expression

DNA methylation phenotypes were generated by dividing the chicken genome into 1000bp windows, yielding a total of 1050176 methylation windows, of which 82426 were located on chromosome Z. The MeDIP-seq reads were mapped to each methylation window and normalised by dividing with the total read count for each individual respectively. Sequencing was performed on an IonProton machine (Thermo Fisher Scientific) using the Torrent Suite software (version 4.4.1) by the National Genomics Infrastructure in Uppsala, Sweden. The sequence depth was on average 3.4X ± 0.97 (standard deviation), the read length was on average 136 ± 15 bp, the raw reads was on average 23.8 million ± 5.2 and the quality score was on average 22 ± 1. The Gene expression dataset has been published previously (Johnsson, Williams et al. 2016) and was based on the NimbleGen 12 × 135K Custom Gene expresson array, mapping to 22628 unique genes composed of Ensembl, RefSeq genes and Expressed Sequence Tags.

### Quantitative Trait Loci (QTL) analysis

Quantitative Trait Loci (QTL) analysis was performed to identify genomic regions associated with the variation found within DNA methylation levels for the 1 million methylation windows. A genetic marker map was generated using 652 SNP markers, of which 542 were fully informative between the original parental animals used to generate the intercross. Average marker distance was ∼16 cM, as per recommendations (Darvasi and Soller 1994). Of these, 36 markers were present on the Z chromosome with a 15cM average marker distance. Note that as the intercross is a linkage-based cross and not a GWAS of an outbred population (which relies on linkage disequilibrium and has built up historical recombinations over hundreds of generations) far fewer markers are required to cover the genome, as it is only required to identify the recombinations that have accrued during the intercrossing (Lynch and Walsh 1998). Details of the genetic marker locations can be found in Johnsson et al (Johnsson, Rubin et al. 2014). Interval mapping was performed using the “qtl2” R-package (Broman, Gatti et al. 2019). This package was used as it is able to correctly analyse sex chromosomes in an advanced intercross. A local (cis) scan was performed, restricted to methylation windows present on the Z chromosome, with the local region considered to be within 50cM up- and down-stream of each methylation window. A trans scan was also performed. In the case of the trans scan, a full genome scan was performed for trans effect methylation QTL that were located on either the autosomes or Z chromosome that affected methylation on the Z chromosome. In addition, a scan was also performed for trans methylation QTL located on the Z chromosome that were associated with methylation present on the autosomes. Sex and batch were set as covariates in the test model, with sex also used as an interactive covariate, where significant (if the LOD score of the sex interaction model was >1 LOD higher than the non-sex interaction model). Significance thresholds were determined via a permutation test with and without sex interactions for both local (cis) and trans methylation QTL. Local (putatively cis) regions were defined as 50cM up and downstream to the closest genetic marker, whilst anything outside this region was defined as trans. For the trans permutations, 20000 random methylation phenotypes were permuted 1000 times each, both for sex and non-sex interaction, and for cis permutations 17000 random phenotypes permuted 1000 times each. From the permutations the top 5 % LOD-scores for each phenotype were saved and from these the top 5% were chosen as significance threshold and the top 20% as the suggestive threshold, respectively. This yielded significance cis LOD-score of 5.73 (sex interaction) and 4.29, (no sex interaction), with suggestive thresholds of 4.87 and 3.58. For the trans thresholds, significance was at LOD-score of 7.70 and 7.68 (sex and non-sex interaction, respectively), whilst the suggestive threshold was 5.92 and 5.93.

Gene expression QTL (eQTL) analysis was performed using R/qtl, using RMA preprocessed (Irizarry, Bolstad et al. 2003) expression levels as quantitative phenotypes with sex and batch as additive covariates. The same criteria for cis-eQTL was applied as for the autosomes (see (Johnsson, Williams et al. 2016), with local eQTL defined as those within +/-50cM of the gene, with trans referring to any other location. Significance thresholds for cis and trans eQTL were 4.0 and 6.0, respectively.

### Male Hyper-Methylated (MHM) region

The MHM region was identified using the transcript deposited in the NCBI GenBank by (Teranishi, Shimada et al. 2001), accession AB046698 (2332 bp), with this being the probe sequence used to identify the region initially. This sequence maps to two genomic locations: chrZ:27375241-27391116 (99.1% match) and chrZ:27329191-27333743 (98.9% match), hereafter referred to as MHMa and MHMb respectively. The MHMa and MHMb regions were corroborated in our dataset and the parameters for methylation levels obtained were used to identify other MHM-like regions. These parameters were: median methylation status per window of > 8.52, a sex difference equal to a Wilcoxon rank sum test/Mann-Whitney test p-value < 1.75e-10 and comprising of five or more adjacent methylation windows (i.e. these values are those identified for the original MHM region in our dataset).

### QTL overlaps

With both mQTL and eQTL mapped it was possible to assess whether any correlations could be found between DNA methylation levels and gene expression which are both associated to a locus. By overlapping the confidence intervals of the mQTL and eQTL, and regressing the gene expression with methylation, genomic regions that putatively control either the methylation or gene expression (or both) were observed. The correlation was tested with all individuals and sex as a factor, and with the sexes separate, yielding 3 models. Any genes that significantly correlated with a methylation window were finally tested for causality using the Network Edge Orientation (NEO) package in R (Aten, Fuller et al. 2008). In this way, it is possible to ascribe hypothetical orientation of the regulatory relationship, whether DNA methylation regulates gene expression or vice versa. Significance using the NEO package is based on the LEO.NB score, which quantifies the support of the best fitting causal model versus the second best fitting model. As both the eQTL and methylation QTL originated from the same genotype (imputed marker position) and thus are treated as a single-marker orientation with a LEO.NB.OCA-score > 1.0 considered significant, and a score of > 0.8 as suggestive.

## Data availability

Gene expression data (generated with Microarray) for the hypothalamus tissue is available at Arrayexpress [https://www.ebi.ac.uk/arrayexpress/experiments/E-MTAB-3154/]. The genotypes scored for the QTL analysis is available at Figshare [https://doi.org/10.6084/m9.figshare.12803876].

The DNA methylation data (generated with MeDIP) is available at:

https://doi.org/10.6084/m9.figshare.12803873

Finally, the readymade QTL cross-file is available at:

## Results

### Dosage Compensation and the Male Hyper Methylated (MHM) Region

To assess the degree of male-biased hyper methylated regions, we first analysed the previously known hyper methylated regions –MHMa and MHMb. These two regions, situated at 27.375Mb and 27.329Mb respectively, had a 3.3 and 3.6-fold increase in methylation in males, respectively, with these ratios being highly significant (max Wilcoxon pvalue < 4e-10 and 1.7e-10, respectively, for each region), see Supplementary Table 1). The original MHM region was hypothesised to be approximately 460kb in length (Teranishi et al. 2001). When we assessed the methylation around these two regions, we find elevated methylation from 27.142Mb-27.40Mb (259-kb long), more accurately demarking this region, see Figure 1. To identify further male biased methylation windows, we performed a chromosome-wide scan calculating the degree of sex bias. Based on the pre-existing MHM region, we then selected all those regions with both a strongly significant sex bias (p<1.75e-10, as compared to the average sex bias in methylation on the Z chromosome being p=0.14 and a 1.69 fold methylation difference between males and females) and with at least five adjacent methylation windows (see methods section).

**Figure 1.**
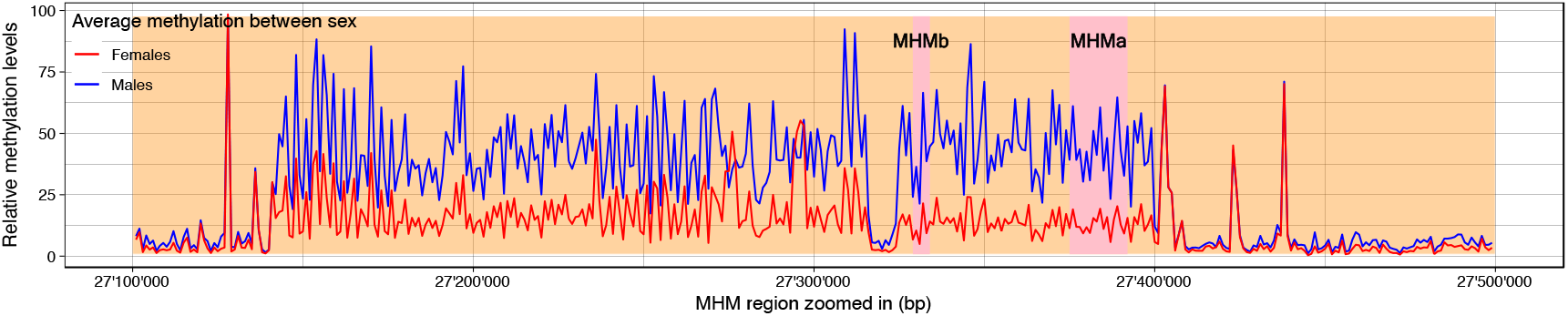
MHMa and MHMb regions (used to identify the original MHM region) and sex differences in methylation levels at these and the surrounding area. Male methylation is shown in blue, female methylation is shown in red.

In total, 19 MHM regions (hereafter referred to as blocks) were identified (see Table 1, Figure 2 and Supplementary Figure 1). Of these continuous blocks, 17 had genes in the local vicinity. In this instance, we defined local as being with 100kb of the MHM block, as in our previous study we found strong correlations between gene expression and DNA methylation even up to 100kb away from the gene itself. To test if dosage compensation acts locally on a gene-by-gene basis or uniformly throughout each block, the methylation levels within these MHM blocks were correlated with the neighbouring genes (see Table 1), i.e. individual methylation windows present within each block were correlated with the expression of adjacent genes, controlling for multiple testing. Of the 17 blocks with adjacent genes, 14 had a significant correlation between at least one methylation window and local gene expression, see Figure 2 and Supplementary Figure 1. Interestingly, neighbouring genes frequently displayed differential correlation with methylation, indicating that these regions seem to be associated with expression on a gene-by-gene basis. In total, 51 unique genes (38 present in our dataset) were adjacent to these MHM-like blocks, with 224 significant correlations with methylation levels (methylation windows) of which 134 correlations were negative and 90 positive (tvalue from linear model). Furthermore, of the 38 genes present in our dataset, 34 had a significant sex bias expression with 20 being expressed higher in males and 14 higher in females (M:F ratio). The average fold difference between males and females on the Z chromosome was 1.22 while for the autosomes this was 1.02. In the case of the original MHM region, apart from the RNAse genes (EST probes X603141644 and X603862378 for the lncRNA *ENSGALG00000051419* in Figure 2) that are almost entirely silent in males, this region (see Figure 2, Supplementary Figure 1, and Table 1) also contains multiple genes that are still male-biased, but below the average degree of male-bias on the Z chromosome. Similarly, these genes tend to be positively correlated with local methylation, where such a correlation exists. This pattern is also replicated in the newly identified MHM regions (see MHM#1 and #2 in Figure 2, and MHM#12,13,14,15,16,19 in Supplementary Figure 1). Therefore, increased methylation in males is associated with a reduction in the differences in male-biased gene expression, but not eliminate it entirely, in both the existing and the new MHM regions. None of the methylation QTL detected on the Z chromosome (either QTL or phenotypes) overlapped with these MHM regions.

**Table 1.** Novel Male Hyper-Methylated (MHM) regions identified in the hypothalamus. The 17 MHM regions containing genes are divided into separate regions, with their location, size, number of probesets present initially given. Also included are the average gene expression values for males and females, the p-value of the sex differences in gene expression, the ratio of male:female gene expression, the number of 1kb windows present within the MHM region that correlate with each gene and the direction of that correlation.

| methbin | chr | pos | avg | median | avg_male | avg_female | pvalue | MF_avg | MF_runavg | FHM_blockid |
| --- | --- | --- | --- | --- | --- | --- | --- | --- | --- | --- |
| chrZ_30195000 | Z | 30195000 | 80.17 | 70.52 | 71.05 | 91.62 | 1.41E-02 | 0.78 | 1.26 | 1 |
| chrZ_30196000 | Z | 30196000 | 52.73 | 48.01 | 42.69 | 65.32 | 2.07E-05 | 0.65 | 1.25 | 1 |
| chrZ_30197000 | Z | 30197000 | 78.65 | 73.65 | 70.17 | 89.28 | 7.32E-03 | 0.79 | 1.23 | 1 |
| chrZ_30198000 | Z | 30198000 | 66.48 | 60.78 | 56.40 | 79.13 | 2.62E-04 | 0.71 | 1.20 | 1 |
| chrZ_30199000 | Z | 30199000 | 69.29 | 61.94 | 61.36 | 79.25 | 1.64E-02 | 0.77 | 1.18 | 1 |
| chrZ_30200000 | Z | 30200000 | 32.70 | 30.17 | 28.75 | 37.65 | 1.92E-02 | 0.76 | 1.16 | 1 |
| chrZ_42633000 | Z | 42633000 | 57.33 | 50.01 | 49.31 | 67.39 | 4.38E-03 | 0.73 | 1.53 | 2 |
| chrZ_42634000 | Z | 42634000 | 81.07 | 76.75 | 67.55 | 98.04 | 1.25E-05 | 0.69 | 1.50 | 2 |
| chrZ_42635000 | Z | 42635000 | 64.55 | 59.29 | 54.35 | 77.36 | 1.73E-04 | 0.70 | 1.46 | 2 |
| chrZ_42636000 | Z | 42636000 | 64.06 | 59.41 | 55.99 | 74.20 | 4.46E-03 | 0.75 | 1.43 | 2 |
| chrZ_42637000 | Z | 42637000 | 87.94 | 82.79 | 77.66 | 100.85 | 1.71E-03 | 0.77 | 1.43 | 2 |
| chrZ_42638000 | Z | 42638000 | 77.38 | 73.39 | 69.06 | 87.82 | 2.81E-02 | 0.79 | 1.42 | 2 |
| chrZ_49068000 | Z | 49068000 | 33.32 | 33.10 | 29.34 | 38.32 | 9.41E-04 | 0.77 | 1.43 | 3 |
| chrZ_49069000 | Z | 49069000 | 103.06 | 93.79 | 88.37 | 121.50 | 2.15E-04 | 0.73 | 1.41 | 3 |
| chrZ_49070000 | Z | 49070000 | 73.53 | 64.76 | 61.14 | 89.06 | 1.55E-04 | 0.69 | 1.38 | 3 |
| chrZ_49071000 | Z | 49071000 | 82.15 | 76.27 | 69.69 | 97.78 | 2.40E-04 | 0.71 | 1.34 | 3 |
| chrZ_49072000 | Z | 49072000 | 109.91 | 100.09 | 91.25 | 133.31 | 1.73E-06 | 0.68 | 1.31 | 3 |
| chrZ_49073000 | Z | 49073000 | 51.46 | 46.02 | 42.84 | 62.27 | 4.95E-04 | 0.69 | 1.27 | 3 |

**Figure 2.**
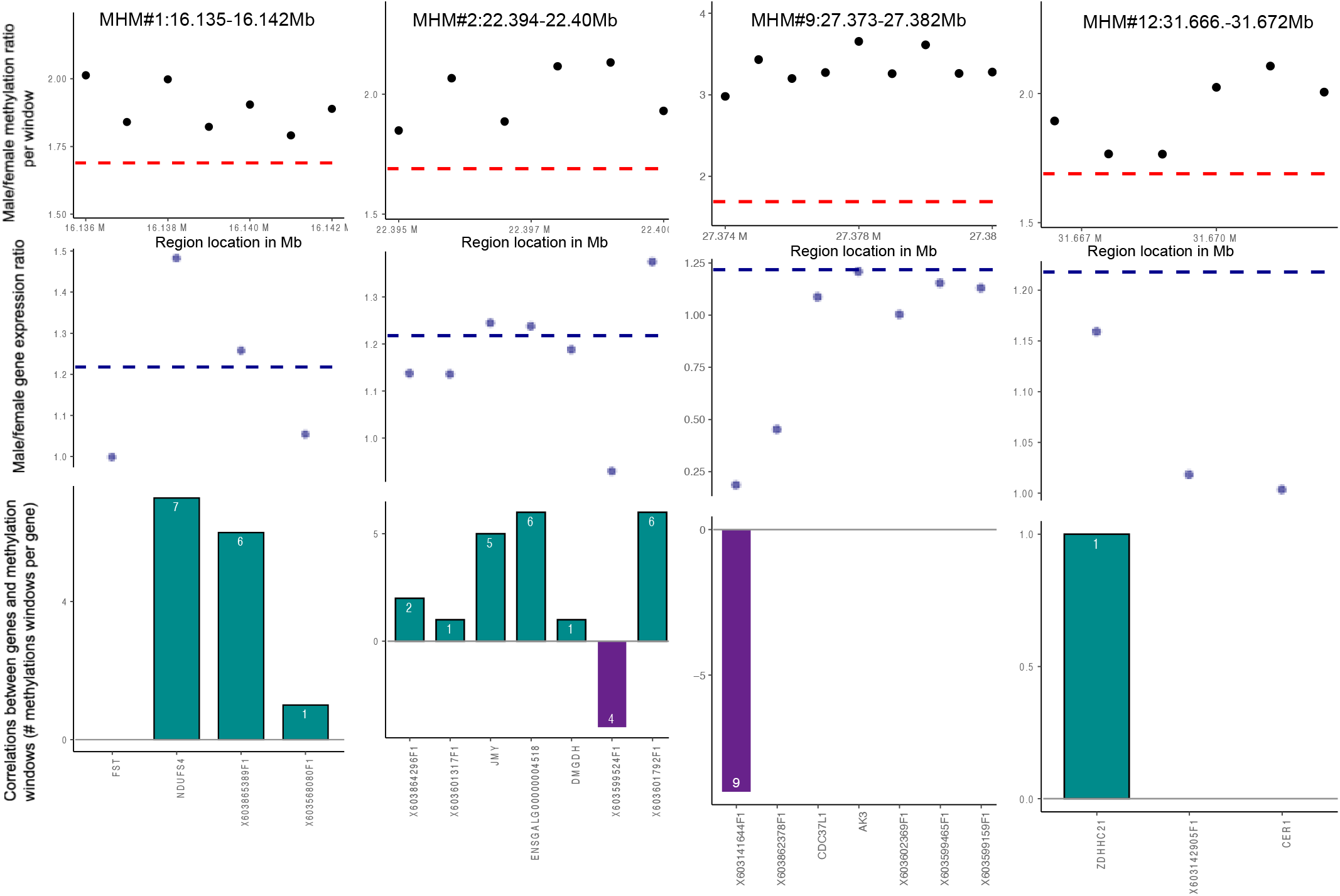
Four of the 19 Novel MHM regions present on the Z chromosome and their effects on gene expression. Panes illustrate regions 1, 2, 9, 12 (selected as being representative of all the regions). Each pane consists of the following: i) The male:female methylation ratio for the 1kb methylation windows that make up the MHM region (each black dot represents the ratio at one methylation window). The red hashed line at the base indicates the average male:female methylation ratio (∼1.7). ii) Male:female gene expression ratio is indicated by the blue dots, one for each gene in the region, with the ratio shown on the left-side y-axis, and the blue hashed line indicating the average male:female gene expression ratio on the Z chromosome (∼1.2). iii) The number of correlations between each gene and the 1kb methylation windows that make up each MHM. The direction of the correlation (positive or negative) is indicated by the bar being above the line (positive, coloured turquoise) or below the line (negative, coloured purple). The number of correlations is indicated on each bar, whilst each gene name is given on the x-axis.

One other MHM region has previously been putatively identified at 73.16-73.173Mb on the Z chromosome by Sun et al. (2019). We also identify this region in our data, though the median methylation threshold fell slightly below the threshold we set, and was therefore excluded initially (i.e. there was a strongly significant sex-difference, but the median level of methylation over all individuals was lower than in the original MHM region). Nevertheless, the region shows very significant DNA methylation levels differences between the sexes (see Supplementary Table 2), with significantly more male DNA methylation. All of the neighbouring genes to these MHM regions were also assessed for potential GO enrichment, with no GO enrichment found for those genes in the immediate vicinity.

### Female Hyper-Methylated Regions

As well as additional MHM regions, a search for regions with a lower than average male: female methylation ratio was also performed to identify regions that showed a relative decrease in DNA methylation in males or an increase in DNA methylation in females. Using a criterion of a significant increase in female methylation, relative to males, we firstly identified a total of 118 1kb windows that were significantly more methylated in females than males (see Table 2 and Supplementary Table 3). Of these, three regions consisted of five or more consecutive female-biased methylated windows. These regions were located at 30195000-30200000bp, 42633000-42638000bp, and 49073000-49073000bp on the Z chromosome. No genes were found in these regions, however. An overlap between methylation QTL and these regions was also performed, though once again no overlaps occurred.

**Table 2.** Novel Female Hyper-Methylated regions identified in the hypothalamus. The position, average and median methylation per window, and the average methylation in males and females per window are all given, as well as the significance of the sex-difference and the average male:female fold ratio.

| id | num_qtl | marker | chr | pos | CI_low_marker | CI_high_marker | CI_low_pos | CI_high_pos | CI_size | num_genes |
| --- | --- | --- | --- | --- | --- | --- | --- | --- | --- | --- |
| hotspot_1 | 11 | Gg_rs14793763 | 1 | 13847380 | Gg_rs13832402 | Gg_rs15194859 | 12488579 | 14847187 | 2358608 | 50 |
| hotspot_2 | 23 | Gg_rs14858437 | 1 | 92741754 | Gg_rs13901810 | Gg_rs13910957 | 91200315 | 101131756 | 9931441 | 129 |
| hotspot_3 | 11 | Gg_rs15060526 | 2 | 8387159 | Gg_rs14132382 | Gg_rs14139143 | 5843745 | 11697701 | 5853956 | 87 |
| hotspot_4 | 64 | Gg_rs15282380 | 3 | 17984384 | X3_16300000 | Gg_rs14327472 | 15489694 | 23999342 | 8509648 | 201 |
| hotspot_5 | 16 | Gg_rs15416272 | 3 | 86515515 | Gg_rs15403420 | Gg_rs15427786 | 79192646 | 91808917 | 12616271 | 177 |
| hotspot_6 | 19 | X4_1267185 | 4 | 1286191 | X4_1267185 | Gg_rs13546113 | 1286191 | 1841819 | 555628 | 39 |
| hotspot_7 | 14 | Gg_rs15679503 | 5 | 22362570 | snp.98.79.91070.S.2 | Gg_rs15685956 | 17614557 | 25665093 | 8050536 | 184 |
| hotspot_8 | 27 | Gg_rs14568888 | 6 | 7742744 | Gg_rs15765462 | Gg_rs15777012 | 6568227 | 10633493 | 4065266 | 72 |
| hotspot_9 | 11 | Gg_rs15828492 | 7 | 2469584 | Gg_rs15826188 | Gg_rs16575534 | 1680380 | 3673817 | 1993437 | 26 |
| hotspot_10 | 18 | Gg_rs13609494 | 12 | 5538321 | Gg_rs13621493 | Gg_rs14974529 | 3076405 | 6304262 | 3227857 | 92 |
| hotspot_11 | 37 | rbl1871 | 14 | 15000631 | Gg_rs15002638 | rbl1871 | 13628710 | 15000631 | 1371921 | 52 |
| hotspot_12 | 12 | Gg_rs13744918 | 17 | 3944254 | Gg_rs15033588 | Gg_rs13744523 | 2473253 | 5465495 | 2992242 | 76 |
| hotspot_13 | 39 | Gg_rs16782623 | Z | 41699011 | Gg_rs16768340 | Gg_rs16114279 | 37374725 | 52128174 | 14753449 | 176 |

### Methylation QTL present on the Z chromosome

Methylation QTL were assessed by performing local (cis) methylation QTL scans restricted solely to the Z chromosome. In addition, trans scans were also performed, where the QTL was located on the Z chromosome, but the target methylation window was free to be present on either the Z chromosome or the autosomes. In total, we identify 18 significant cis methylation QTL and 53 significant trans methylation QTL that are based on the Z chromosome, with a further 20 suggestive cis methylation QTL and 528 trans methylation QTL. As expected, most of the methylation QTL (n=528) had a significant sex interaction effect. This is expected due to the large differences in Z chromosome methylation between males and females, with males possessing two methylated chromosomes (ZZ) and females only one (ZW). A full list of all methylation QTL can be found in Supplementary Table 4. In addition, 51 expression QTL (eQTL) were identified on the Z chromosome (either as a QTL or the trans-effect phenotype of a QTL), see Supplementary Table 5. ready.

### Trans Methylation QTL Hotspots Affecting the Z chromosome

To identify trans-acting hotspots, we identified where multiple methylation QTL were associated with the same marker and had overlapping confidence intervals. Of the 619 methylation QTL on the Z chromosome, these mapped to 141 different SNP loci. Of these loci, 13 were associated with multiple methylation windows/phenotypes (10 or more methylation windows associated with each marker, respectively). These hotspots on average spanned 5.87Mb of physical distance in the genome (found by taking the shared overlapping confidence intervals and finding the minimum overlapping size), see Table 3. Of note, all bar one (n=12) of these trans hotspots were located on the autosomes, but regulated variation in methylation on the Z chromosome. Of these 12, 3 were previously identified as regulating methylation variation on the autosomes in this intercross (Höglund et al. 2020), on chromosomes 3 (at 18Mb, hotspot 4), 6 (at 7.7Mb, hotspot 6) and 7 (at 2.4 Mb, hotspot 9). One hotspot was located on the Z chromosome (at 41.7Mb, hotspot 13, with this hotspot spread over three adjacent SNPs, rs16768340, rs16782623, rs14016786, see Supplementary Table 4) regulated variation in methylation on different windows in the Z chromosome, as well as some methylation windows on the autosomes. Thus, whilst the majority of regulation in methylation variation appears to be located on the autosomes, with these loci then regulating methylation on the Z chromosome, there is also some regulation of methylation variation by the Z chromosome itself, and even a small amount of autosomal regulation from the Z chromosome. The genes present within these hotspots were further checked for potential enrichment via gene ontology analysis, using DAVID 6.8 (https://david.ncifcrf.gov/). In total 3 hotspots showed enrichment using the DAVID 6.8 database: the hotspot (ID#2) at chr1@91.7MB contained genes enriched for immunoglobulin-fold/domain, the hotspot (ID#5 in Table 3) on chr3@86.5Mb had genes enriched for the activity of glutathione and metabolism of cytochrome P450, and the hotspot (ID#6 in table 3) on chr4@1.3Mb contained genes enriched for activity with rhodopsin, see Supplementary Table 6. The hotspots and their distribution across the genome are illustrated in Figure 3. Gene enrichment analysis was also performed for the target genomic regions in the vicinity (±10kb) of each methylation window associated with a methylation QTL hotspot. Some enrichment was found for hotspot ID#4 (located on chromosome 3 at 17.98Mb), however, this result was non-significant (Bonferroni *p*-value > 0.05).

**Table 3.**
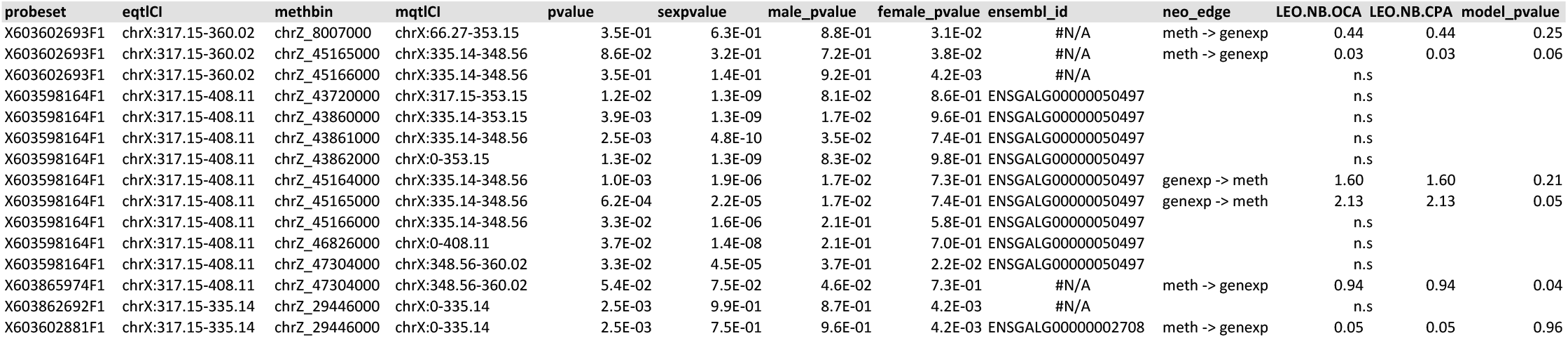
List of trans methylation QTL hotspots. Table shows the number of methylation QTL present for each hotspot, its chromosome and base-pair position (nearest marker), and the confidence interval of each hotspot. The number of genes present within the intervals as determined by ensembl.org is also given.

**Figure 3.**
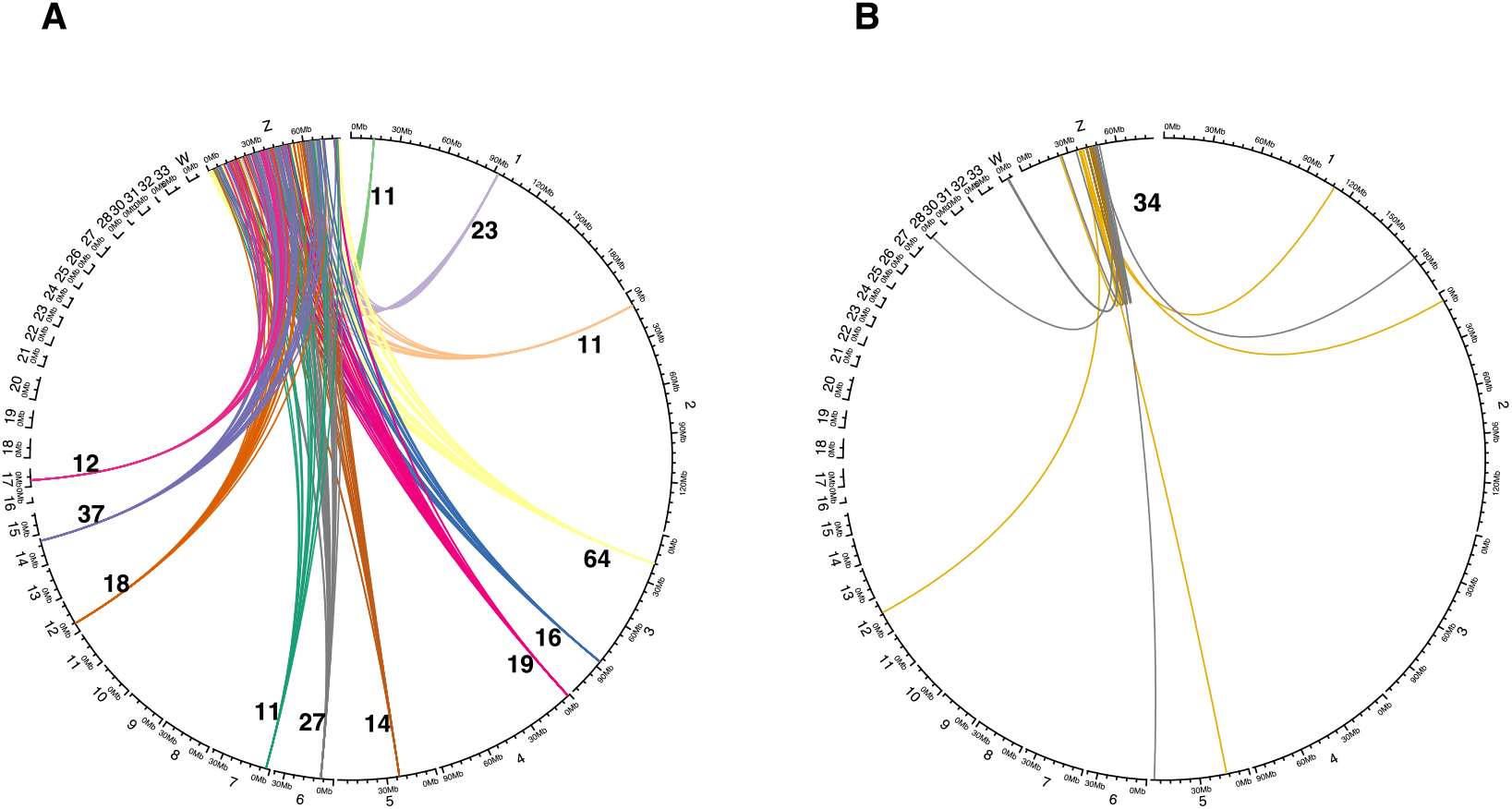
Circle plot showing the location of trans methylation QTL hotspots that affect DNA methylation variation on the Z chromosome. (A) The 12 autosomal hotspots affecting Z DNA methylation, and (B) the single Z chromosome hotspot affecting Z and autosomal methylation.

### Causality Analysis Between Methylation and Gene Expression on the Z chromosome

In total 360 overlaps were found between eQTL and methylation QTL. These were methylation and expression QTL where either the QTL or methylation phenotype were located on the Z chromosome. The overlapping phenotypes (gene expression and methylation) were tested for association using a linear model. Of these, 15 overlaps were significant after applying an FDR-based multiple testing corrections. Eleven of the overlaps were significant (p-value < 0.05, FDR corrected) using all individuals, while 3 were significant (p-value < 0.05, FDR corrected) using only females, and 1 was significant (p-value < 0.05, FDR corrected) using only males, see Table 4. These overlaps contained 5 unique probesets belonging to 2 unique genes and 3 ESTs. The gene LINGO1 (*ENSGALG00000002708*; chr10:3212741-3290778) is an immunoglobulin domain protein (Yang, Jiang et al. 2022). Immunoglobulin activity was also found in the methylation QTL hotspot on chromosome 3. Additionally, the 15 overlaps were tested with NEO, a network edge orientated method which uses the underlying QTL genotype as anchors for the network (Aten et al., 2008), to assess the orientation of the observed correlation. Four of the overlaps had a LEO.NB.OCA-score > 0.3. Both eQTL and mQTL originated from the same genotype (imputed marker position) and thus are treated as a single-marker orientation where a LEO.NB.OCA-score > 1.0 is significant. Hence, our results indicate that the EST X603598164F1 (gene id: *ENSGALG00000050497, chrZ:44706094-44707218*) influences the methylation levels in the region of chrZ:45163000-45165000, see Table 4. This gene has been retired on the GalGal6 genome, with no known function. In addition one further EST (X603865974) was suggestive (LEO.NB.OCA >0.8), with methylation appearing to drive gene expression in this case. However, as the model p-value was significant, this means that other models (gene expression driving methylation) cannot be ruled out.

## DISCUSSION

Using this wild x domestic paradigm to analyse DNA methylation and its regulation on the Z chromosome in the chicken, we firstly identify over 600 methylation QTL that affect methylation on the Z chromosome. Of these, the majority of trans effect loci are located on the autosomes but affecting the Z chromosome. There were also examples of the reverse, with trans methylation QTL deriving from the Z chromosome but affecting DNA methylation on the autosomes. Furthermore, these trans methylation QTL were concentrated into a small number of hotspots located on the autosomes (n=12), with one hotspot also present on the Z chromosome itself, associated with methylation on the Z chromosome and the autosomes, respectively. A total of five genes on the Z chromosome were also candidates for causality between gene expression and methylation, with two passing the network-edge-based threshold for significance. Of these, one appears to be a retired gene, whilst the other is an EST of no known function, with the former indicating gene expression affects methylation, whilst the latter indicates that methylation was modifying gene expression.

The nature of the intercross (a wild bird intercrossed with domestics) allows us to identify consistent differences in methylation that exist between wild and domestic chickens and the regions that associate with and potentially regulate them. With regards to the methylation QTL hotspots identified, it is noteworthy that these are almost all based on the autosomes, with only one situated on the Z chromosome itself. Therefore, the regulation of domestication-based phenotypes with loci present on the Z chromosome appears to generally be autosomally regulated, although the reverse (where autosomal gene expression is regulated by the sex chromosomes) also occurs. Interestingly, three of the hotspots previously identified as regulating DNA methylation in domestication (primarily via reducing DNA methylation in domestic birds) also appear to regulate DNA methylation on the Z chromosome (Höglund 2020).

As well as the regulation of variation in methylation, we also identified additional Male Hyper-Methylated regions present on the Z chromosome. Unlike the initial MHM region found (Teranishi, Shimada et al. 2001), which identified that the lncRNAs present were completely switched off in males, the regions we identify appear to instead decrease male gene expression, though rather than reduce it entirely, it is instead down-regulated to levels more closely found in females (i.e. reduced male gene expression, relative to female gene expression). This is despite the regions having a similar pattern of sex-differentiated methylation as is seen in the original MHM region. Further, the strength of the methylation differences between sexes was greater in the new regions we identified when compared to the region at 74Mb (although we also identify the 74Mb MHM region as well). These genes thus appear to be linked to sex-based differences between males and females. No methylation QTL overlap these regions, implying that these regions are not responsible for regulating variation in methylation, which would then fit with these regions instead regulating more basal sex-differences rather than between-population variation. This idea is reinforced when considering the functions of the genes in these regions.

Of the 18 known genes that are present within the MHM regions, their functions can be broadly divided into learning/behaviour, bone allocation, development, reproduction, growth/metabolism and methyl transferase activities. These tie-in well with the known sex-differences that exist in the chicken. Starting with behaviour, strong behavioural differences exist between males and female chickens (Vallortigara, Cailotto et al. 1990, Nätt, Agnvall et al. 2014, Elfwing, Nätt et al. 2015, Bélteky, Agnvall et al. 2018). In particular, females have decreased anxiety-related behaviour, though this may be test-dependent (Schutz, Kerje et al. 2002, Campler, Jöngren et al. 2009, Johnsson, Williams et al. 2016, Johnsson, Henriksen et al. 2018, Fogelholm, Inkabi et al. 2019). Of the genes present in the MHM, four are related to behaviour or neurogenesis. The gene *SLC1A1* has been shown to play a role in obsessive compulsive disorder and sterotype behaviour (Zike, Chohan et al. 2017, Huang, Liu et al. 2021), as well as schizophrenia susceptibility (Horiuchi, Iida et al. 2012, Li, Su et al. 2020). Anxiety behaviour in chickens has previously been shown to be related to schizophrenia, depression and other mood-based disorders in humans, even sharing some of the same susceptibility loci (Johnsson, Williams et al. 2016, Johnsson, Henriksen et al. 2018). Furthermore, the OCD effects arising from *SLC1A1* are stronger in males, so sex-differences in the gene effects have already been demonstrated (Wendland, Moya et al. 2009, Veenstra-VanderWeele, Xu et al. 2012). ZDHHC2I is a major palmitoyl acyltransferase, with decreasing expression leading to increased depression-like behaviours (Gorinski, Bijata et al. 2019). *Homer1* also has functions relating to learning and memory (Clifton, Cameron et al. 2017), and also causes susceptibility to Alzheimers (Urdánoz-Casado, Sánchez-Ruiz de Gordoa et al. 2021). In the case of the latter, these effects are strongly sex-dependent, only occurring in women.

Continuing with bone allocation, female chickens have a complex bone allocation, whereby during egg production the hard outer cortical bone is first mobilised into soft, spongy medullary bone in the centre of the femur, before then being transferred to create the egg shell (one of the major limiting factors in egg production) (Bloom, Domm et al. 1958, Mueller, Schraer et al. 1964). Therefore male and female chickens differ markedly in their bone metabolism – males possess almost no medullary bone, whilst female medullary bone deposition is strongly associated with reproductive output (Johnsson, Gustafson et al. 2012, Johnsson, Rubin et al. 2014, Johnsson, Jonsson et al. 2015). Of the genes in the MHM, *Homer1* has numerous beneficial effects in osteoblasts including beta-catenin stabilization (Rybchyn, Brennan-Speranza et al. 2021). *FST* (foillistatin) is also a powerful regulator of bone metabolism (Gajos-Michniewicz, Piastowska et al. 2010). *CER1* has also been found to regulate bone mineral density and be associated with fracture risk. Of note, these effects are found to be strongest in post-menopausal women(Koromila, Dailiana et al. 2012, Koromila, Georgoulias et al. 2013). *TLE4* is a critical mediator of osteoblasts and *runx2-*dependent bone development in the mouse (Shin, Theodorou et al. 2021). Finally, *NANS* affects skeletal development in zebrafish knock-outs (van Karnebeek, Bonafé et al. 2016). Continuing with reproduction-related genes, the gene *FST* plays a critical role in mouse uterine receptivity and decidualization (Fullerton, Monsivais et al. 2017), whilst the gene *JMY* mediates spermatogenesis in mice (Liu, Fan et al. 2020) as well as asymmetric division and cytokinesis in mouse oocytes (Sun, Sun et al. 2011). Finally, *CLTA4* is involved in the maintenance of chronic inflammation in endometriosis and infertility (Abramiuk, Bębnowska et al. 2021).

The final category of genes present in the MHM regions affected growth and metabolism, whilst two methyltransferase genes were also present. Large differences in growth and bodyweight exist in the chicken, with males often twice the bodyweight of females. The gene *FST*, as well as affecting reproduction-related phenotypes, also leads to increased muscle weight in mice when over-expressed (Iyer, Chugh et al. 2021). *DMGDH* affects body growth through insulin-like growth factor (Baker et al 1993), whilst also affecting selenium status in pregnant women (Mao, Vanderlelie et al. 2016). *ABHD3* regulates adipogenic differentiation and lipid metabolism (Linke, Overmyer et al. 2020). Finally, *BHMT* is a methyltransferase, as is *DMGDH*.

In the current study, we have restricted the investigation to a single tissue, albeit repeated in 124 individuals. As such, we are confident that these MHM regions and methylation QTL are present in this tissue type. However the ubiquity of these methylated regions in other tissues must be verified. This opens up the possibility that multiple further MHM regions also exist, but are only present in specific tissue types. This could allow fine-scale regulation of sex differences in a tissue-specific manner. We also assess female-specific hyper-methylated regions, but these were found to be very sparse and had very few genes present, suggesting these are of less importance. In summary, by using a large number of replicates that are assessed for all methylated loci on the Z chromosome, we identify both novel MHM regions in an intra-specific/inter-population framework, as well as the role that domestication plays in the regulation of the Z chromosome and the genes on it.

## Acknowledgements

The research was carried out within the framework of the Swedish Centre of Excellence in Animal Welfare Science, and the Linköping University Neuro-network. SNP genotyping was performed by the Uppsala Sequencing Center. Bioinformatic support was provided by NBIS (National Bioinformatics Infrastructure Sweden). The project was supported by grants from the European Research Council (Advanced grant GENEWELL and Consolidator grant FERALGEN 772874), the Swedish Research Council (VR), Carl Tryggers Stiftelse, and the Linköping University Neuro-network.

## Table Legends

Table 4. NEO causality of gene regulation of methylation. The probeset and the methylation window being tested, along with their confidence interval is presented. In addition, the genotype p-value, the sex p-value (also broken down into male and female), as well as the actual causality statistics (leo.nb.oca and cpa and the model p-value) are all shown.

Supplementary Table 1. MHMa and MHMb regions. The position, average and median methylation per window, and the average methylation in males and females per window are all given, as well as the significance of the sex-difference and the average male:female fold ratio.

Supplementary Table 2. The previously identified MHM region at 73Mb. The position, average and median methylation per window, and the average methylation in males and females per window are all given, as well as the p-value and significance of the sex-difference and the average male:female fold ratio. Note for the significance of the sex difference, these are classified as non-significant, significant (including a multiple testing correction), and significant at the same level as the original MHM region.

Supplementary Table 3. All female hyper-methylated regions. The three FHM blocks (continuous regions) are highlighted in orange and indicated with their block ID in a separate column. The position, average and median methylation per window, and the average methylation in males and females per window are all given, as well as the significance of the sex-difference and the average male:female fold ratio.

Supplementary Table 4. List of local (cis) and trans methylation QTL present on the Z chromosome. The phenotype of each methylation QTL (methylation window), nearest marker to the methylation QTL, LOD score, confidence interval, and nearest marker to each confidence are given, as well as whether the QTL is cis or trans in effect, are all given.

Supplementary Table 5. Expression QTL (eQTL) present on the Z chromosome. Closest marker, LOD score, confidence interval, presence or absence of sex interaction, and nearest marker to the confidence interval are all presented.

Supplementary Table 6. GO enrichments from methylation QTL hotspots. Category, GO term, p-value (absolute and also FDR controlled), genes involved and fold enrichment are all given.

## REFERENCES

Abramiuk, M., D. Bębnowska, R. Hrynkiewicz, P. Niedźwiedzka-Rystwej, G. Polak, J. Kotarski, J. Roliński and E. Grywalska (2021). “CLTA-4 expression is associated with the maintenance of chronic inflammation in endometriosis and infertility.” Cells 10(3): 487.

Aten, J. E., T. F. Fuller, A. J. Lusis and S. Horvath (2008). “Using genetic markers to orient the edges in quantitative trait networks: the NEO software.” BMC Systems Biology 2(1): 34.

Bélteky, J., B. Agnvall, L. Bektic, A. Höglund, P. Jensen and C. Guerrero-Bosagna (2018). “Epigenetics and early domestication: differences in hypothalamic DNA methylation between red junglefowl divergently selected for high or low fear of humans.” Genetics Selection Evolution 50(1): 13.

Bloom, M. A., L. V. Domm, A. V. Nalbandov and W. Bloom (1958). “Medullary bone of laying chickens.” American Journal of Anatomy 102(3): 411–453.

Broman, K. W., D. M. Gatti, P. Simecek, N. A. Furlotte, P. Prins, Ś. Sen, B. S. Yandell and G. A. Churchill (2019). “R/qtl2: Software for Mapping Quantitative Trait Loci with High-Dimensional Data and Multiparent Populations.” Genetics 211(2): 495–502.

Campler, M., M. Jöngren and P. Jensen (2009). “Fearfulness in red junglefowl and domesticated White Leghorn chickens.” Behavioural Processes 81(1): 39–43.

Clifton, N. E., D. Cameron, S. Trent, L. H. Sykes, K. L. Thomas and J. Hall (2017). “Hippocampal Regulation of Postsynaptic Density Homer1 by Associative Learning.” Neural Plast 2017: 5959182.

Darvasi, A. and M. Soller (1994). “Optimum spacing of genetic markers for determining linkage between marker loci and quantitative trait loci.” Theoretical and Applied Genetics 89(2-3): 351–357.

Elfwing, M., D. Nätt, V. C. Goerlich-Jansson, M. Persson, J. Hjelm and P. Jensen (2015). “Early stress causes sex-specific, life-long changes in behaviour, levels of gonadal hormones, and gene expression in chickens.” PLoS One 10(5): e0125808.

Ellegren, H., L. Hultin-Rosenberg, B. Brunström, L. Dencker, K. Kultima and B. Scholz (2007). “Faced with inequality: chicken do not have a general dosage compensation of sex –linked genes.” BMC biology 5(1): 1–12.

Fang, H., C. M. Disteche and J. B. Berletch (2019). “X inactivation and escape: pepigenetic and structural features.” Frontiers in cell and developmental biology 7: 219.

Fogelholm, J., S. Inkabi, A. Höglund, R. Abbey-Lee, M. Johnsson, P. Jensen, R. Henriksen and D. Wright (2019). “Genetical Genomics of Tonic Immobility in the Chicken.” Genes 10(5): 341.

Fullerton, P. T., Jr., D. Monsivais, R. Kommagani and M. M. Matzuk (2017). “Follistatin is critical for mouse uterine receptivity and decidualization.” Proc Natl Acad Sci U S A 114(24): E4772–e4781.

Gajos-Michniewicz, A., A. W. Piastowska, J. A. Russell and T. Ochedalski (2010). “Follistatin as a potent regulator of bone metabolism.” Biomarkers 15(7): 563–574.

Gaston, K. and M. Fried (1995). “CpG methylation has differential effects on the binding of YY1 and ETS proteins to the bi-directional promoter of the Surf-1 and Surf-2 genes.” Nucleic acids research 23(6): 901–909.

Gorinski, N., M. Bijata, S. Prasad, A. Wirth, D. Abdel Galil, A. Zeug, D. Bazovkina, E. Kondaurova, E. Kulikova, T. Ilchibaeva, M. Zareba-Koziol, F. Papaleo, D. Scheggia, G. Kochlamazashvili, A. Dityatev, I. Smyth, A. Krzystyniak, J. Wlodarczyk, D. W. Richter, T. Strekalova, S. Sigrist, C. Bang, L. Hobuß, J. Fiedler, T. Thum, V. S. Naumenko, G. Pandey and E. Ponimaskin (2019). “Attenuated palmitoylation of serotonin receptor 5-HT1A affects receptor function and contributes to depression-like behaviors.” Nat Commun 10(1): 3924.

Guerrero-Bosagna, C. (2019). From epigenotype to new genotypes: pRelevance of epigenetic mechanisms in the emergence of genomic evolutionary novelty. Seminars in cell &developmental biology, Elsevier.

Heyn, H., S. Moran, I. Hernando-Herraez, S. Sayols, A. Gomez, J. Sandoval, D. Monk, K. Hata, T. Marques-Bonet and L. Wang (2013). “DNA methylation contributes to natural human variation.” Genome research 23(9): 1363–1372.

Höglund, A., R. Henriksen, J. Fogelholm, A. M. Churcher, C. G. Guerrero-Bosagna, A. Martinez Barrio, M. Johnsson, P. Jensen and D. Wright (2020). “The methylation landscape and its role in domestication and gene regulation in the chicken.” Nature Ecology and Evolution 4(12): 1713–1724.

Horiuchi, Y., S. Iida, M. Koga, H. Ishiguro, Y. Iijima, T. Inada, Y. Watanabe, T. Someya, H. Ujike, N. Iwata, N. Ozaki, H. Kunugi, M. Tochigi, M. Itokawa, M. Arai, K. Niizato, S. Iritani, A. Kakita, H. Takahashi, H. Nawa and T. Arinami (2012). “Association of SNPs linked to increased expression of SLC1A1 with schizophrenia.” Am J Med Genet B Neuropsychiatr Genet 159b(1): 30–37.

Huang, X., J. Liu, J. Cong and X. Zhang (2021). “Association Between the SLC1A1 Glutamate Transporter Gene and Obsessive-Compulsive Disorder in the Chinese Han Population.” Neuropsychiatr Dis Treat 17: 347–354.

Irizarry, R. A., B. M. Bolstad, F. Collin, L. M. Cope, B. Hobbs and T. P. Speed (2003). “Summaries of Affymetrix GeneChip probe level data.” Nucleic Acids Research 31(4): e15.

Itoh, Y., E. Melamed, X. Yang, K. Kampf, S. Wang, N. Yehya, A. Van Nas, K. Replogle, M. R. Band and D. F. Clayton (2007). “Dosage compensation is less effective in birds than in mammals.” Journal of biology 6(1): 1–15.

Iyer, C. C., D. Chugh, P. J. Bobbili, A. J. B. Iii, A. E. Crum, A. F. Yi, B. K. Kaspar, K. C. Meyer, A. H. M. Burghes and W. D. Arnold (2021). “Follistatin-induced muscle hypertrophy in aged mice improves neuromuscular junction innervation and function.” Neurobiol Aging 104: 32–41.

Johnsson, M., I. Gustafson, C.-J. Rubin, A.-S. Sahlqvist, K. B. Jonsson, S. Kerje, O. Ekwall, O. Kämpe, L. Andersson, P. Jensen and D. Wright (2012). “A Sexual Ornament in Chickens Is Affected by Pleiotropic Alleles at HAO1 and BMP2, Selected during Domestication.” PLoS Genetics 8(8): e1002914.

Johnsson, M., R. Henriksen, J. Fogelholm, A. Höglund, P. Jensen and D. Wright (2018). “Genetics and genomics of social behavior in a chicken model.” Genetics 209(1): 209–221.

Johnsson, M., K. B. Jonsson, L. Andersson, P. Jensen and D. Wright (2015). “Genetic Regulation of Bone Metabolism in the Chicken: pSimilarities and Differences to Mammalian Systems.” PLoS Genetics 11(5): e1005250.

Johnsson, M., C. J. Rubin, A. Höglund, A. S. Sahlqvist, K. Jonsson, S. Kerje, O. Ekwall, O. Kämpe, L. Andersson, P. Jensen and D. Wright (2014). “The role of pleiotropy and linkage in genes affecting a sexual ornament and bone allocation in the chicken.” Molecular ecology 23(9): 2275–2286.

Johnsson, M., M. J. Williams, P. Jensen and D. Wright (2016). “Genetical genomics of behavior: pa novel chicken genomic model for anxiety behavior.” Genetics 202(1): 327–340.

Kasowski, M., S. Kyriazopoulou-Panagiotopoulou, F. Grubert, J. B. Zaugg, A. Kundaje, Y. Liu, A. P. Boyle, Q. C. Zhang, F. Zakharia and D. V. Spacek (2013). “Extensive variation in chromatin states across humans.” Science 342(6159): 750–752.

Kilpinen, H., S. M. Waszak, A. R. Gschwind, S. K. Raghav, R. M. Witwicki, A. Orioli, E. Migliavacca, M. Wiederkehr, M. Gutierrez-Arcelus and N. I. Panousis (2013). “Coordinated effects of sequence variation on DNA binding, chromatin structure, and transcription.” Science 342(6159): 744–747.

Koromila, T., Z. Dailiana, S. Samara, C. Chassanidis, C. Tzavara, G. P. Patrinos, V. Aleporou-Marinou and P. Kollia (2012). “Novel sequence variations in the CER1 gene are strongly associated with low bone mineral density and risk of osteoporotic fracture in postmenopausal women.” Calcif Tissue Int 91(1): 15–23.

Koromila, T., P. Georgoulias, Z. Dailiana, E. E. Ntzani, S. Samara, C. Chassanidis, V. Aleporou-Marinou and P. Kollia (2013). “CER1 gene variations associated with bone mineral density, bone markers, and early menopause in postmenopausal women.” Hum Genomics 7(1): 21.

Li, W., X. Su, T. Chen, Z. Li, Y. Yang, L. Zhang, Q. Liu, M. Shao, Y. Zhang, M. Ding, Y. Lu, H. Yu, X. Fan, M. Song and L. Lv (2020). “Solute Carrier Family 1 (SLC1A1) Contributes to Susceptibility and Psychopathology Symptoms of Schizophrenia in the Han Chinese Population.” Front Psychiatry 11: 559210.

Linke, V., K. A. Overmyer, I. J. Miller, D. R. Brademan, P. D. Hutchins, E. A. Trujillo, T. R. Reddy, J. D. Russell, E. M. Cushing, K. L. Schueler, D. S. Stapleton, M. E. Rabaglia, M. P. Keller, D. M. Gatti, G. R. Keele, D. Pham, K. W. Broman, G. A. Churchill, A. D. Attie and J. J. Coon (2020). “A large-scale genome-lipid association map guides lipid identification.” Nat Metab 2(10): 1149–1162.

Liu, Y., J. Fan, Y. Yan, X. Dang, R. Zhao, Y. Xu and Z. Ding (2020). “JMY expression by Sertoli cells contributes to mediating spermatogenesis in mice.” Febs j 287(24): 5478–5497.

Lynch, M. and B. Walsh (1998). Genetics and Analysis of Quantitative traits. Sunderland, MA, Sinauer Associates.

Mank, J. and H. Ellegren (2009). “All dosage compensation is local: gene–by-gene regulation of sex-biased expression on the chicken Z chromosome.” Heredity 102(3): 312–320.

Mank, J. E. (2013). “Sex chromosome dosage compensation: pdefinitely not for everyone.” Trends in genetics 29(12): 677–683.

Mann, I. K., R. Chatterjee, J. Zhao, X. He, M. T. Weirauch, T. R. Hughes and C. Vinson (2013). “CG methylated microarrays identify a novel methylated sequence bound by the CEBPB| ATF4 heterodimer that is active in vivo.” Genome research 23(6): 988–997.

Mao, J., J. J. Vanderlelie, A. V. Perkins, C. W. Redman, K. R. Ahmadi and M. P. Rayman (2016). “Genetic polymorphisms that affect selenium status and response to selenium supplementation in United Kingdom pregnant women.” Am J Clin Nutr 103(1): 100–106.

McVicker, G., B. van de Geijn, J. F. Degner, C. E. Cain, N. E. Banovich, A. Raj, N. Lewellen, M. Myrthil, Y. Gilad and J. K. Pritchard (2013). “Identification of genetic variants that affect histone modifications in human cells.” Science 342(6159): 747–749.

Melamed, E. and A. P. Arnold (2007). “Regional differences in dosage compensation on the chicken Z chromosome.” Genome biology 8(9): 1–10.

Mueller, W. J., R. Schraer and H. Scharer (1964). “Calcium metabolism and skeletal dynamics of laying pullets.” The Journal of nutrition 84(1): 20–26.

Nalvarte, I., J. Rüegg and C. Guerrero-Bosagna (2018). Intrinsic and Extrinsic Factors That Influence Epigenetics. Epigenetics and Assisted Reproduction, CRC Press: 99–114.

Nätt, D., B. Agnvall and P. Jensen (2014). “Large sex differences in chicken behavior and brain gene expression coincide with few differences in promoter DNA-methylation.” PLoS One 9(4): e96376.

Rybchyn, M. S., T. C. Brennan-Speranza, D. Mor, Z. Cheng, W. Chang, A. D. Conigrave and R. S. Mason (2021). “The mTORC2 Regulator Homer1 Modulates Protein Levels and Sub-Cellular Localization of the CaSR in Osteoblast-Lineage Cells.” Int J Mol Sci 22(12).

Schutz, K., S. Kerje, O. Carlborg, L. Jacobsson, L. Andersson and P. Jensen (2002). “QTL analysis of a red junglefowl x White Leghorn intercross reveals trade-off in resource allocation between behavior and production traits.” Behavior Genetics 32(6): 423–433.

Shin, T. H., E. Theodorou, C. Holland, R. Yamin, C. L. Raggio, P. F. Giampietro and D. A. Sweetser (2021). “TLE4 Is a Critical Mediator of Osteoblast and Runx2-Dependent Bone Development.” Front Cell Dev Biol 9: 671029.

Sun, D., D. L. Maney, T. S. Layman, P. Chatterjee and V. Y. Soojin (2019). “Regional epigenetic differentiation of the Z Chromosome between sexes in a female heterogametic system.” Genome research 29(10): 1673–1684.

Sun, S. C., Q. Y. Sun and N. H. Kim (2011). “JMY is required for asymmetric division and cytokinesis in mouse oocytes.” Mol Hum Reprod 17(5): 296–304.

Teranishi, M., Y. Shimada, T. Hori, O. Nakabayashi, T. Kikuchi, T. Macleod, R. Pym, B. Sheldon, I. Solovei and H. Macgregor (2001). “Transcripts of the MHM region on the chicken Z chromosome accumulate as non-coding RNA in the nucleus of female cells adjacent to the DMRT1 locus.” Chromosome Research 9(2): 147–165.

Urdánoz-Casado, A. J. Sánchez-Ruiz de Gordoa, M. Robles, B. Acha, M. Roldan, M. V. Zelaya, I. Blanco-Luquin and M. Mendioroz (2021). “Gender-Dependent Deregulation of Linear and Circular RNA Variants of HOMER1 in the Entorhinal Cortex of Alzheimer’s Disease.” Int J Mol Sci 22(17).

Vallortigara, G., M. Cailotto and M. Zanforlin (1990). “Sex differences in social reinstatement motivation of the domestic chick (Gallus gallus) revealed by runway tests with social and nonsocial reinforcement.” Journal of Comparative Psychology 104(4): 361.

van Karnebeek, C. D., L. Bonafé, X. Y. Wen, M. Tarailo-Graovac, S. Balzano, B. Royer-Bertrand, A. Ashikov, L. Garavelli, I. Mammi, L. Turolla, C. Breen, D. Donnai, V. Cormier-Daire, D. Heron, G. Nishimura, S. Uchikawa, B. Campos-Xavier, A. Rossi, T. Hennet, K. Brand-Arzamendi, J. Rozmus, K. Harshman, B. J. Stevenson, E. Girardi, G. Superti-Furga, T. Dewan, A. Collingridge, J. Halparin, C. J. Ross, M. I. Van Allen, A. Rossi, U. F. Engelke, L. A. Kluijtmans, E. van der Heeft, H. Renkema, A. de Brouwer, K. Huijben, F. Zijlstra, T. Heise, T. Boltje, W. W. Wasserman, C. Rivolta, S. Unger, D. J. Lefeber, R. A. Wevers and A. Superti-Furga (2016). “NANS-mediated synthesis of sialic acid is required for brain and skeletal development.” Nat Genet 48(7): 777–784.

Veenstra-VanderWeele, J., T. Xu, A. M. Ruggiero, L. R. Anderson, S. T. Jones, J. A. Himle, J. L. Kennedy, M. A. Richter, G. L. Hanna and P. D. Arnold (2012). “Functional studies and rare variant screening of SLC1A1/EAAC1 in males with obsessive-compulsive disorder.” Psychiatr Genet 22(5): 256–260.

Vicoso, B. and D. Bachtrog (2011). “Lack of global dosage compensation in Schistosoma mansoni, a female-heterogametic parasite.” Genome Biology and Evolution 3: 230–235.

Wendland, J. R., P. R. Moya, K. R. Timpano, A. P. Anavitarte, M. R. Kruse, M. G. Wheaton, R. F. Ren-Patterson and D. L. Murphy (2009). “A haplotype containing quantitative trait loci for SLC1A1 gene expression and its association with obsessive-compulsive disorder.” Arch Gen Psychiatry 66(4): 408–416.

Yang, H., L. Jiang, Y. Zhang, X. Liang, J. Tang, Q. He, Y. M. Luo, C. N. Zhou, L. Zhu, S. S. Zhang, K. Xiao, P. L. Zhu, J. Wang, Y. Li, F. L. Chao and Y. Tang (2022). “Anti-LINGO-1 antibody treatment alleviates cognitive deficits and promotes maturation of oligodendrocytes in the hippocampus of APP/PS1 mice.” J Comp Neurol 530(10): 1606–1621.

Yin, Y., E. Morgunova, A. Jolma, E. Kaasinen, B. Sahu, S. Khund-Sayeed, P. K. Das, T. Kivioja, K. Dave and F. Zhong (2017). “Impact of cytosine methylation on DNA binding specificities of human transcription factors.” Science 356(6337): eaaj2239.

Zhang, M., F.-B. Yan, F. Li, K.-R. Jiang, D.-H. Li, R.-L. Han, Z.-J. Li, R.-R. Jiang, X.-J. Liu and X.-T. Kang (2017). “Genome-wide DNA methylation profiles reveal novel candidate genes associated with meat quality at different age stages in hens.” Scientific Reports 7(1): 1–15.

Zike, I. D., M. O. Chohan, J. M. Kopelman, E. N. Krasnow, D. Flicker, K. M. Nautiyal, M. Bubser, C. Kellendonk, C. K. Jones, G. Stanwood, K. F. Tanaka, H. Moore, S. E. Ahmari and J. Veenstra-VanderWeele (2017). “OCD candidate gene SLC1A1/EAAT3 impacts basal ganglia-mediated activity and stereotypic behavior.” Proc Natl Acad Sci U S A 114(22): 5719–5724.

